# Modeling of dynamic cerebrovascular reactivity to spontaneous and externally induced CO2 fluctuations in the human brain using BOLD-fMRI

**DOI:** 10.1101/176891

**Authors:** Prokopis C. Prokopiou, Kyle T. S. Pattinson, Richard G. Wise, Georgios D. Mitsis

## Abstract

In this work, we investigate the regional characteristics of the dynamic interactions between arterial CO2 and BOLD (dynamic cerebrovascular reactivity - dCVR) during normal breathing and hypercapnic, externally induced step CO2 challenges. To obtain CVR curves at each voxel, we use a custom set of basis functions based on the Laguerre and gamma basis sets. This allows us to obtain robust dCVR estimates both in larger regions of interest (ROIs) as well as in individual voxels. We also implement classification schemes to identify brain regions with similar dCVR characteristics. Our results reveal considerable variability of dCVR across different brain regions, as well as during different experimental conditions (normal breathing and hypercapnia), suggesting a differential response of cerebral vasculature to spontaneous CO2 fluctuations and larger, externally induced CO2 changes. The clustering results suggest that anatomically distinct brain regions exhibit different dCVR curves that do not exhibit the standard, positive valued curves that have been previously reported using measurements of global cerebral blood flow or using larger ROIs. They also revealed a consistent set of dCVR cluster shapes for resting and forcing conditions that exhibited different distribution patterns across brain voxels.

## 1. Introduction

Cerebral blood flow (CBF) is regulated by multifactorial homeostatic mechanisms that maintain its value relatively constant. The ability of the brain to achieve this in response to changes in perfusion pressure is termed cerebral autoregulation (Lucas et al. 2010; Mitsis et al. 2002, 2004; Panerai 1998; Tzeng and Ainslie 2014). In addition to perfusion pressure, the cerebrovascular bed is highly responsive to local tissue metabolism (Attwell et al. 2010; Iadecola and Nedergaard 2007) and arterial levels of carbon dioxide (CO2) (Battisti-Charbonney, Fisher, and Duffin 2011; Brugniaux et al. 2007; Duffin 2011; Ratnatunga and Adiseshiah 1990). The CBF response to arterial CO2 changes is termed cerebrovascular reactivity (CVR) and has been assessed using both transcranial Doppler ultrasound (Mitsis et al. 2004; Poulin, Liang, and Robbins 1996) and functional magnetic resonance imaging (fMRI) (Tancredi and Hoge 2013; Wise et al. 2004; Yezhuvath et al. 2009). Furthermore, the important role of CVR in cerebral autoregulation has been suggested (Mitsis et al. 2004; Tzeng et al. 2014).

To assess CVR, spontaneous physiological variability (Golestani et al. 2015; Mitsis et al. 2004; Prokopiou et al. 2016; Wise et al. 2004), arterial gas manipulation protocols such as end-tidal forcing and prospective control (Blockley et al. 2011; Pattinson et al. 2009; Slessarev et al. 2007; Wise et al. 2007) or, more recently, sinusoidally modulated gas stimuli (Blockley et al 2017), as well as controlled breathing (Bright and Murphy 2013; Murphy, Harris, and Wise 2011) have been used. The advantages of CO2 as a vasoactive stimulus have been suggested (Fierstra et al. 2013). Also, spontaneous activity is a desirable stimulus as it removes the need for any external interventions, making it applicable to all populations.

Many studies have implicated CVR as a potential marker for the prognosis and evaluation of disorders related to cerebrovascular dysfunction. Such disorders include arterial stenosis (Mandell, Han, Poublanc, Crawley, Stainsby, et al. 2008), enhanced risk of stroke (Gur, Bova, and Bornstein 1996; Markus and Cullinane 2001; Silvestrini et al. 2000), and other steno-occlusive diseases such as Moyamoya disease (Donahue et al. 2013; Mikulis et al. 2005). CVR has also been associated with small-vessel diseases (Conklin et al. 2010, 2011) and has been shown to correlate with seizure susceptibility in epileptic patients with brain arteriovenous malformation (Fierstra et al. 2011). In addition, studies have illustrated that CVR evaluation could be applied to identify patients with Alzheimer’s disease (Marmarelis et al. 2013, 2016), and Alzheimer’s disease patients who are at higher risk of rapid cognitive decline (Silvestrini et al. 2011). Additional important applications of CVR include calibration of the fMRI signal (Hoge 2012), detection of abnormal neurovascular coupling and physiological effects of pharmacological agents (Chen and Parrish 2009; Hayen et al. 2017), and brain tumor presurgical planning (Zaca, Hua, and Pillai 2011).

When PaCO2 changes with respect to normocapnia, assuming that oxygen consumption remains constant, the blood-oxygen-level-dependent signal obtained with functional magnetic resonance imaging (BOLD–fMRI) can be used as a surrogate for changes in regional CBF (Fierstra et al. 2013). This enables the acquisition of time-courses with a high spatial resolution that account for the sensitivity of the cerebrovascular bed to contemporaneous changes in PaCO2, and allows for investigation of the variability of CVR in different regions of the brain.

The vast majority of the BOLD-based CVR studies define and quantify CVR as the slope of the percent change in the BOLD signal and the CO2 stimulus (Robbins, Swanson, and Howson 1982). While the largest portion of the literature deals with regions of interest (ROIs) defined in the gray matter (GM) where the signal-to-noise ratio (SNR) is high (Bokkers et al. 2010; Bright and Murphy 2013; Wise et al. 2004; Yezhuvath et al. 2009), a few studies have investigated CVR in the brain white matter (WM) (Thomas et al. 2014) and ventricles (Thomas et al. 2013). These studies illustrated that CVR in the brain WM is positive but significantly lower than in the GM, and that cerebrospinal fluid (CSF)-rich regions in the brain, such as the lateral ventricles, exhibit an apparently negative BOLD-CVR.

Recent studies have also addressed the dynamic interactions between hypercapnic, externally induced step CO2 challenges and the BOLD signal, i.e. dynamic CVR (dCVR) (Duffin et al. 2015; Poublanc et al. 2015). The response delay observed between CO2 and BOLD was associated to the time constant of a linear monoexponential system, which corresponds to the dCVR that describes the dynamic interactions between these two physiological signals. In this framework, the time constant of the monoexponential system was estimated in a voxel-wise manner, for a group of patients with known steno-occlusive disease. The estimated response delay at each voxel was then used to identify regions with reduced vasodilatory reserve, associated with the disease pathophysiology (Sobczyk et al. 2014).

In the present study, we investigate in detail the regional characteristics of dCVR using spontaneous and hypercapnic step changes in CO2 (end-tidal forcing) using BOLD-fMRI. We initially estimate dCVR curves within larger functionally and anatomically predefined ROIs, including the brainstem respiratory control centers in the human brain. We use both linear and nonlinear models based on Laguerre function expansions and we show that the effects of CO2 on the BOLD signal are predominantly linear for both experimental conditions. Subsequently, we investigate the regional variability of dCVR in a voxel-wise fashion. To achieve this, we construct a custom basis set based on Laguerre and gamma functions to achieve more robust estimation, and we estimate voxel-specific dCVR curves. We subsequently use the results to construct maps of key dCVR curve features such as total area, peak value, time-to-peak, and power, for each experimental condition, and we use the dCVR feature maps to perform statistical comparisons between the two experimental conditions. Finally, we perform clustering analysis on the estimated voxel-specific dCVR curves in order to identify brain regions with similar dCVR characteristics. Our results suggest that it is possible to obtain reliable dCVR estimates from spontaneous fluctuations using the proposed methodology. The spontaneous and forcing dCVR curves overall exhibit similar characteristics; however, regionally specific differences that are protocol-specific are also revealed. Finally, the clustering analysis suggests the existence of several different dCVR shapes with considerably different characteristics that are correlated to different major brain anatomical structures.

## 2. Methods

### 2.1. Experimental methods

This work is an extended analysis of the experimental data presented in (Pattinson et al. 2009). 12 right-handed healthy volunteers, aged 32 ± 5 years (3 female) participated in this study after giving written informed consent in accordance with the Oxfordshire Clinical Research Ethics committee.

#### 2.1.1. Respiratory protocol

During scanning sessions, subjects were fitted with a facemask (Hans Rudolph, Kansas City, MO, USA) attached to a breathing system, which delivered mixtures of air, O2, and CO2. Continuous recordings of tidal CO2 and O2 (CD-3A and S-3A; AEI Technologies, Pittsburgh, PA, USA), respiratory volume (VMM-400, Interface Associates, Laguna Niguel, CA, USA) and oxygen saturations (9500 Multigas Monitor, MR Equipment Corp., NY, USA), were acquired. It was assumed that the end-tidal partial pressure of CO2 (P_ET_CO2) is a suitable surrogate for PaCO2, and therefore, P_ET_CO2 can be used as the stimulus for CBF.

The study was divided into two parts. For the first part of the study, the subjects were asked to perform no particular task, other than to remain awake with their eyes open and breathe air (resting-state experiment). In the second part of the study, P_ET_CO2 and P_ET_O2 were targeted using dynamic end-tidal forcing (DEF) (Robbins et al. 1982). The CO2 challenges were delivered via a computer controlled gas mixing system (Wise et al. 2007). The CO2 challenges were designed to raise the subjects’ P_ET_CO2 by either 2 or 4 mmHg above a baseline level maintained at 1 mmHg above their natural P_ET_CO2. Representative P_ET_CO2 time series during both conditions are shown in Fig. 1.

**Fig. 1.**
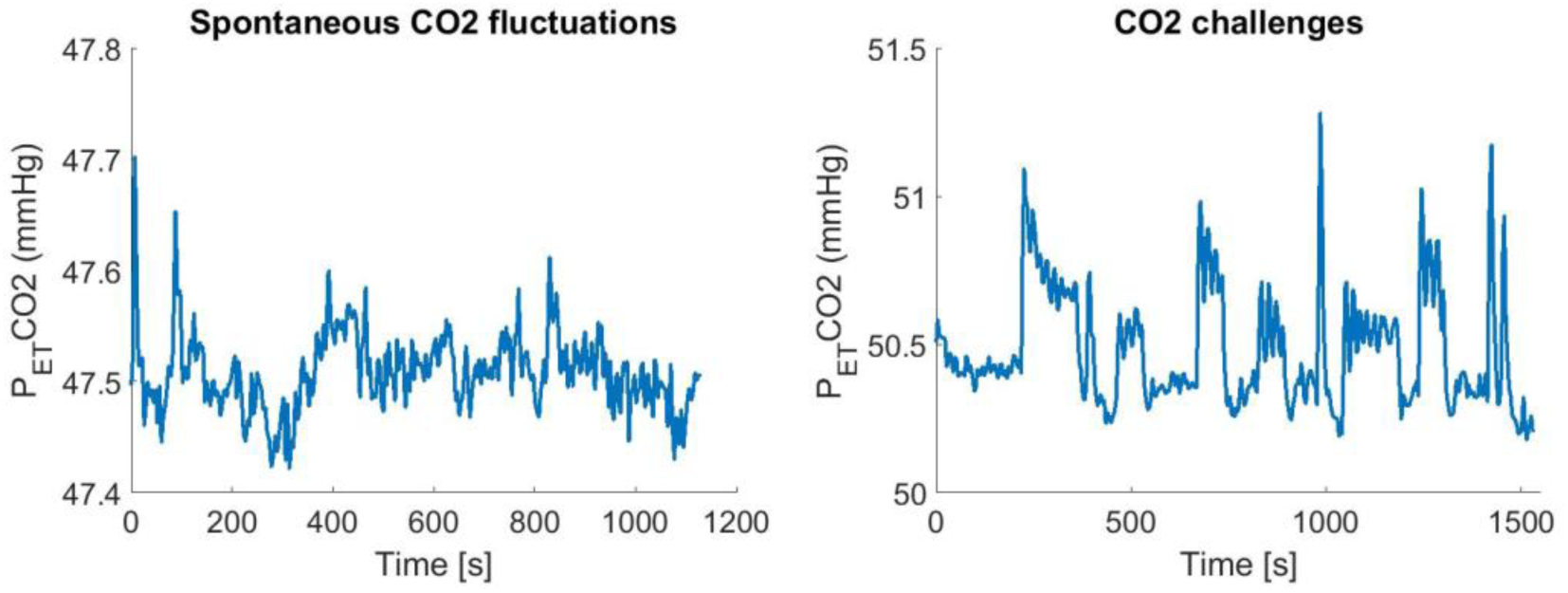
Example of changes in P_ET_CO2 in one representative subject. (left panel) Spontaneous fluctuations during the resting state experiment (right panel) Externally induced step CO2 challenges delivered via a gas control system (dynamic end-tidal forcing).

#### 2.1.2. BOLD imaging

Two thousand seven hundred T2* weighted echo planar imaging (EPI) volumes were acquired on a Siemens Trio 3T scanner. The field of view comprised 16 coronal oblique slices of the brainstem (sequence parameters: *TE = 30 ms*, *TR =1 s*, voxel size 2.5 × 2.5 × 3 *mm*^3^, flip angle 70°).

Although the study was divided in two parts, scanning was continuous. The first 1130 images (18 minutes, 50 seconds) comprised the normal breathing (resting state) experiment. The duration of the first part of the study was determined based upon (Wise et al. 2004), but was prolonged to account for the lower SNR in the brainstem. The final 1530 images (25 minutes, 30 seconds) comprised the CO2 stimulation experiment. The duration of the second part was determined by adaptation of a similar CO2 challenge protocol (Pedersen, Fatemian, and Robbins 1999) for use in the MRI scanner. A high resolution T1-weighted structural scan (voxel size 1 × 1 × 1 *mm*^3^) was also acquired to aid registration to a common stereotactic space of reference.

### 2.2. Data analysis

#### 2.2.1. Data preprocessing

The basic pre-statistical analysis of the data was carried out using FSL (FMRIB, Oxford, UK (Jenkinson et al. 2012)), as has been previously described in (Pattinson et al. 2009). In brief, pre-processing of the BOLD images included spatial smoothing by using a Gaussian kernel of 3.5*mm* FWHM, high-pass temporal filtering, motion correction, registration with T1-weighted anatomic images, and normalization to the Montreal Neurological Institute (MNI)-152 template space, with resolution of 2 × 2 × 2 *mm*^3^. Furthermore, anatomical and functional ROIs were obtained in the MNI space, corresponding to anatomical areas within the area scanned (pons, medulla, thalamus, and putamen) and to areas that revealed increased activity in response to the hypercapnic CO2 challenges (rostral dorsal pons (Kölliker-Fuse / parabrachial nucleus), inferior pons nuclei (ventral respiratory group), the left ventral posterior lateral nucleus of the thalamus, and the left ventrolateral and bilateral ventroanterior nuclei of the thalamus).

The recorded P_ET_CO2 time series were shifted by 3 seconds, to account for the time it takes for the blood to travel from the lungs to the brain tissue, and for the vessels in the tissue to react to the change in CO2 concentration (Liu et al. 2012; Poulin et al. 1996).

#### 2.2.2. Mathematical modelling

Dynamic CO2 reactivity was assessed using linear (impulse response) and non-linear (Volterra kernel) models. In this context, we employed the discrete time Volterra Model (DVM) for a *Q*-th order non-linear system, which is given by

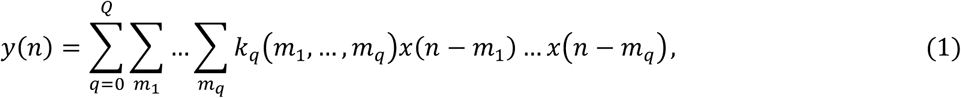

where *y*(*n*) denotes the output (i.e. %BOLD change) and *x*(*n*) the input (i.e. P_ET_CO2 change) of the system at time *n*, respectively, *k*_*q*_ (*m*_*1*_*, …, m*_*q*_) denotes the *q*-th order Volterra kernel of the system, and *Q*denotes the model order.

When *Q* = 1, the right-hand side of (1) reduces to the convolution between the input and the first order Volterra kernel, *k*_1_(*m*_1_), which corresponds to the impulse response of a linear system describing the linear effect of the past input values on the output. Similarly, when *Q* = 2, in addition to the linear term, the right-hand side of (1) consists of a nonlinear term that corresponds to the nonlinear second-order convolution between the input and the second order Volterra kernel, *k*_2_(*m*_1_, *m*_2_), which describes the effect of pairwise interactions (products) of past input values on the output.

The Volterra kernels can be estimated efficiently from the input-output data using a functional expansion technique in terms of an orthonormal basis set (Marmarelis 1993), which is given by

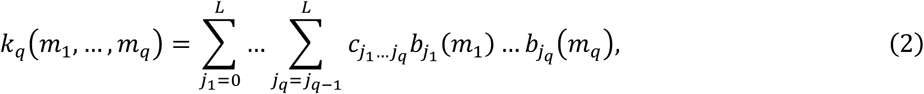

where {*b*_*j*_(*m*);*j* = 0, …, *L*;*m* = 0, …, *M*} is a set of *L*+1 orthonormal basis functions, *c*_*j*_ is the unknown expansion coefficient of the *j*-th order basis function, and *M* the memory of the system. Combining (1) and (2), the DVM can be re-expressed in a compact matrix form as

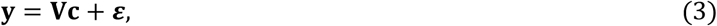

where ***V*** denotes a matrix the values of which are convolutions of the input with the basis functions. The vector ***c*** of the unknown expansion coefficients can be estimated using ordinary least squares

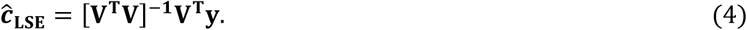

A critical issue arising in the application of the functional expansion technique is the proper choice of the basis set, as it may considerably influence the final estimates. In this work, dCVR was initially investigated within large ROIs using the first (*Q*= 1) and second (*Q*= 2) order DVM, where the unknown values of the Volterra kernels were estimated by employing a set of Laguerre basis functions. The Laguerre basis has been extensively used in the literature, particularly in the case of physiological systems, as they constitute a complete set in [0 ∞) and they exhibit exponentially decaying behavior, which makes them a suitable choice for modeling causal, finite-memory systems (Marmarelis 2004). The *j*-th order discrete time Laguerre function is given by

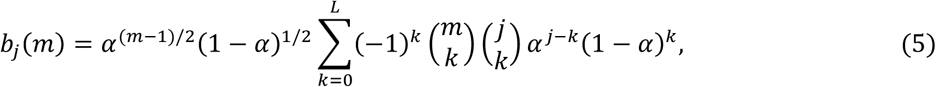

where *α* (0 < *α* < 1) is a parameter that determines the rate of exponential decline of these functions, with larger values corresponding to slower decay.

The values for the model order (*Q*) and number of Laguerre functions (*L*) used in the model, and the parameter *α* were selected based on model performance, which was assessed in terms of the normalized mean squared error (NMSE) between the measured output (i.e. %BOLD change) and the model prediction given by (1). To prevent overfitting, particularly in the case of normal breathing (resting state) BOLD measurements where the SNR is considerably lower, the range for *L* and *α* were selected to be **2** < ***L*** < 6, and 0 < *α* < 0.6 respectively. The comparison of the NMSE values suggested that the dynamic relation between CO2 and BOLD is mainly linear (i.e. *Q* = 1), for both experimental conditions (*p*-values are shown in Table 1). Therefore, in the following, we present results obtained using linear (impulse response) dynamic models.

**Table 1.** The *p*-values corresponding to the statistical comparison between the NMSE values achieved by a linear (*Q*=1) and non-linear (*Q*=2) DVM, in different ROIs. The statistical comparisons were performed using the Kruskal-Wallis nonparametric one-way ANOVA test.

| BRAIN REGION | Dynamic end-tidal forcing | Normal Breathing |
| --- | --- | --- |
| Kölliker-Fuse / parabrachial group | 0.64 | 0.86 |
| Anteroventral thalamic nucleus | 0.33 | 0.49 |
| Ventrolateral thalamic nucleus | 0.53 | 0.65 |
| Ventral Posterior lateral thalamic nucleus | 0.49 | 0.81 |
| Ventral Respiratory Group | 0.36 | 0.60 |
| Retrotrapezoid nucleus | 0.69 | 0.91 |

Our main purpose was to estimate dCVR curves at single voxels, where the SNR is lower. To this end, we constructed a custom, reduced basis set based on gamma density functions, which have been widely used to model the hemodynamic response function (HRF) (Friston et al. 1998). We considered gamma pdfs as described in (Hossein-Zadeh, Ardekani, and Soltanian-Zadeh 2003; Knuth, Ardekani, and Helpern 2001) given by

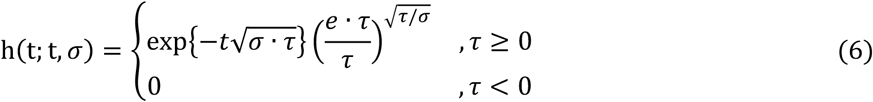

where *τ* and *σ* determine the location of the peak and width, respectively. Guided by the range of linear (impulse response) dynamics between P_ET_CO2 and BOLD that were initially estimated in the larger ROIs using the Laguerre basis functions, we constructed an extended set of gamma functions by varying *τ* and *σ* to span the entire range of the CVR dynamics observed in different brain regions (Fig. 2 – top panel). Subsequently, we applied singular value decomposition (SVD) on this extended set to obtain a reduced set of orthonormal functions that account for the major fraction of the variability in this set. The results yielded two singular vectors (Fig. 2 – bottom panel), as it was found that the two absolutely largest singular values accounted for more than 90% of the extended set variability.

**Fig. 2.**
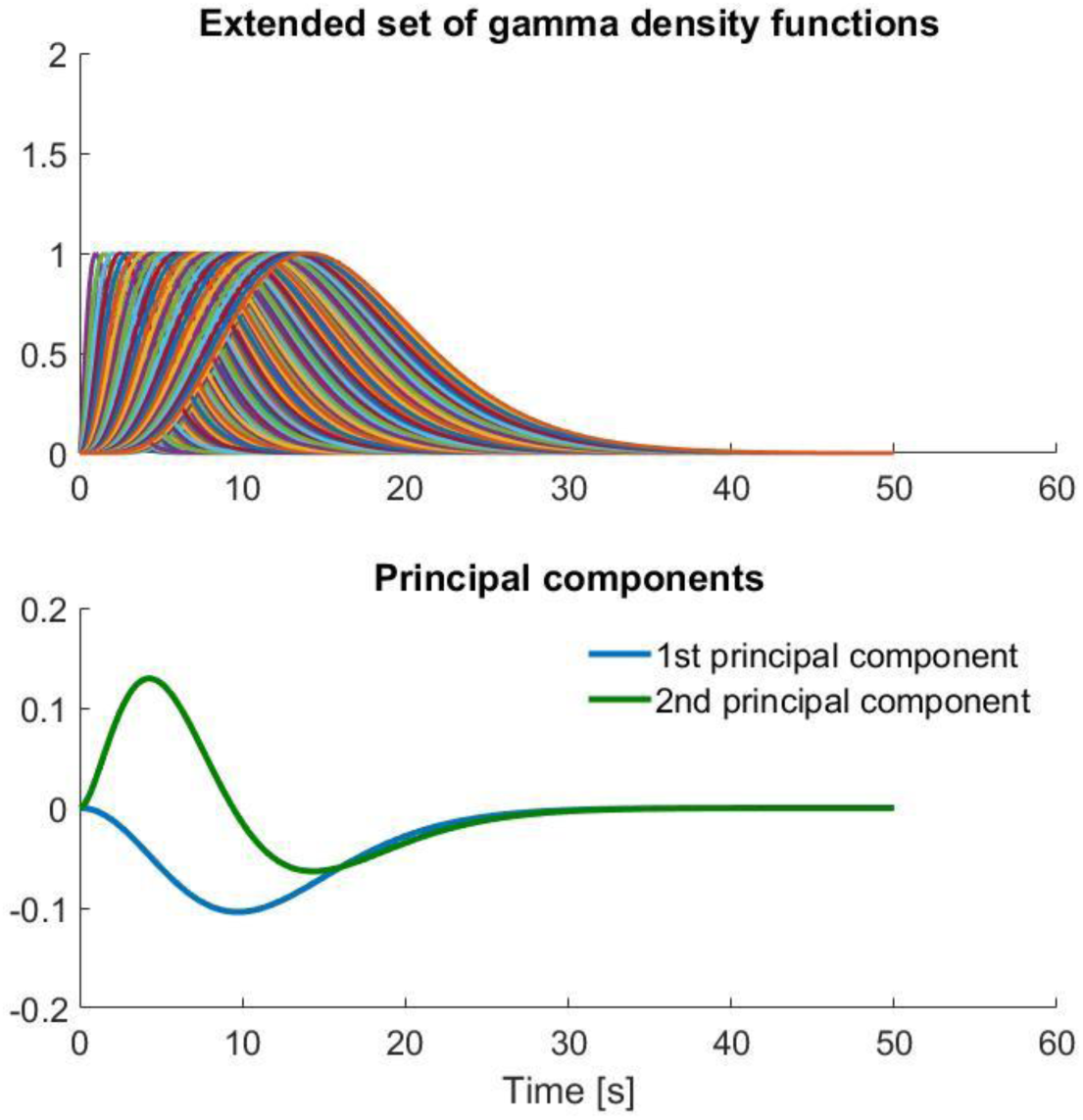
(top panel) Extended set of gamma basis functions. The location of the peak and the memory of each function were varied in accordance to the dCVR curves obtained with the Laguerre basis in large functionally and anatomically defined ROIs. (bottom panel) Reduced set of orthonormal functions, which account for 90% of the variance of the extended set, produced using singular value decomposition. The two orthonormal functions forming the reduced set were used as basis functions in (2) for modeling dCVR.

For both the ROI and voxel-specific analyses, the dCVR curve estimates were obtained using equations (1)-(4) along with the set of two functions of Fig. 2 (bottom panel). For the voxel-specific analysis, we constructed maps of the key features of the voxel-specific dCVR, such as area, peak and time-to-peak values, which illustrate the variability of dCVR across the brain. The area of the dCVR curve corresponds to the steady state CVR value that is typically used as an index of CO2 reactivity in the literature (e.g. Yezhuvath et al. 2009). The peak value describes the maximum instantaneous CO2 reactivity. The power corresponds to the dCVR curve sum-of-squares, and the time-to-peak corresponds to the time lag of the maximum instantaneous CO2 reactivity and may be used to assess how fast a particular voxel/ROI responds to CO2 changes.

In addition, we also performed cluster analysis on the shape of the voxel-wise dCVRs, using unsupervised clustering (k-means) along with the silhouette criterion for the evaluation of clustering performance (Kaufman et al. 2005; Rousseeuw 1987). To perform clustering, the values of the dCVR estimates were normalized to a unit energy function with respect to the sum of squares of all time points (Orban et al. 2014).

## 3. Results

### 3.1.1. ROI analysis

Table 1 illustrates the *p*-values of the Kruskal-Wallis nonparametric one-way ANOVA test between the NMSE values from all subjects achieved by linear and non-linear models, using the DVM with *Q*= 1, and *Q*= 2, respectively, for different ROIs. Both models were identified using the functional expansion technique along with the Laguerre basis. In this context, the null hypothesis was that the NMSE values obtained from both models originate from the same distribution. The *p*-values suggest that we could not reject the null hypothesis, implying that the dynamic relation between P_ET_CO2 and %BOLD is predominantly linear for both experimental conditions.

Fig. 3 shows representative dCVR curves within different ROIs using the basis functions obtained after applying SVD on the extended set of gamma pdfs (Fig. 2 – bottom panel). The initial part of the dCVR curves suggests a similar response to spontaneous and externally induced larger CO2 changes; however, the curves corresponding to resting conditions exhibited a more pronounced late undershoot, which is largely absent from the forcing curves (see also the voxel-wise results below). Representative model output predictions achieved using the gamma and Laguerre basis sets for the left anteroventral nucleus of the thalamus functional ROI under end-tidal forcing and normal breathing (resting state) conditions are shown in Fig. 4. The gamma and Laguerre models yielded similar model predictions, explaining a large fraction of the slower variations in the BOLD signal during end-tidal forcing and normal breathing.

**Fig. 3.**
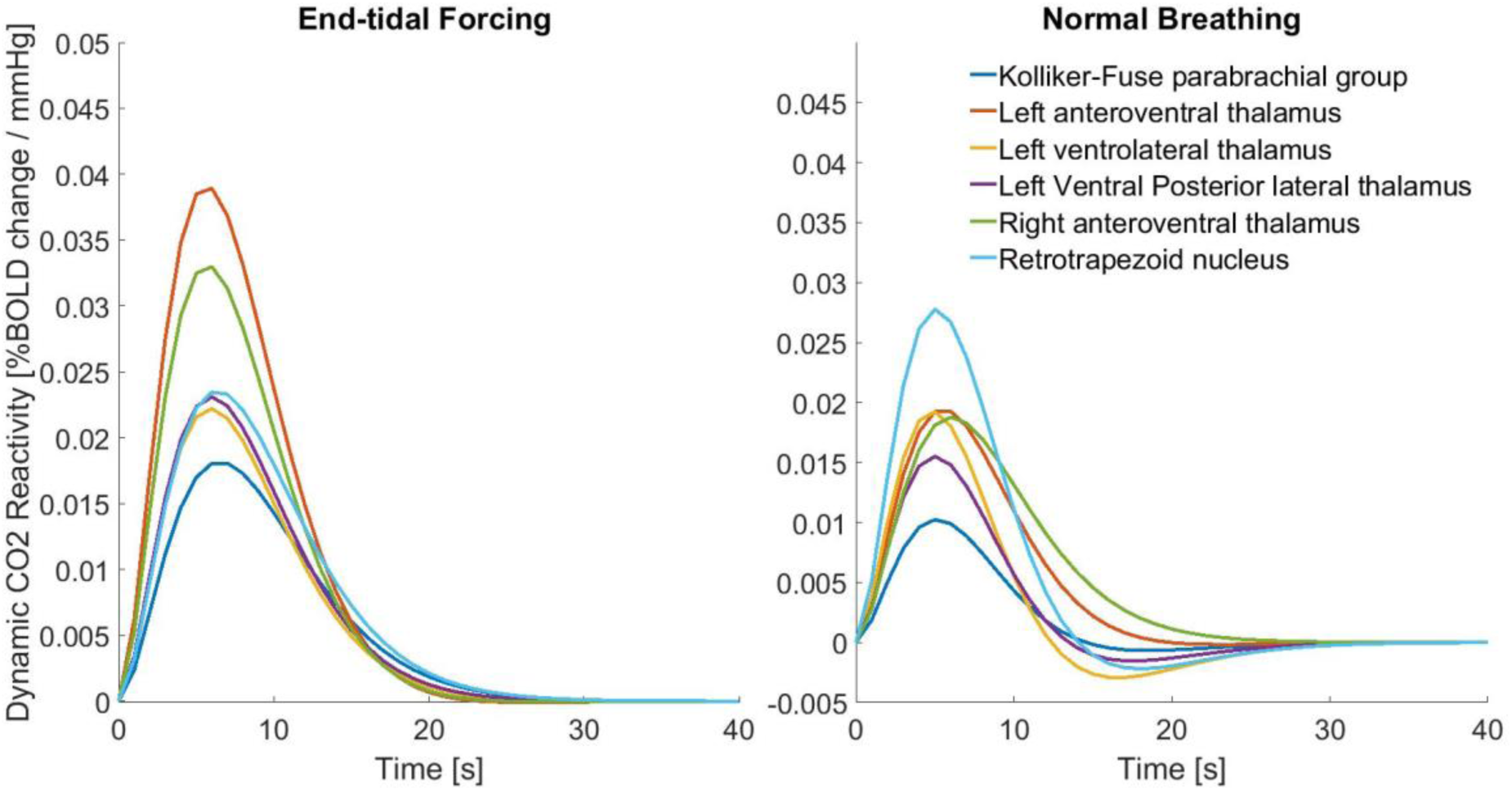
Dynamic cerebrovascular reactivity (dCVR) in different ROIs during forcing (left panel) and resting (right panel) conditions, obtained using the reduced gamma function basis set (Fig. 2 – bottom panel). The regional variability of dCVR is evident. The undershoot observed during normal breathing is absent during forcing conditions. AV: anteroventral thalamus VL: ventrolateral thalamus, VRG: Ventral Respiratory Group, KF/PB: Kolliker-Fuse parabrachial group, VPL: Ventral Posterior lateral thalamus, RTN: Retrotrapezoid nucleus.

**Fig. 4.**
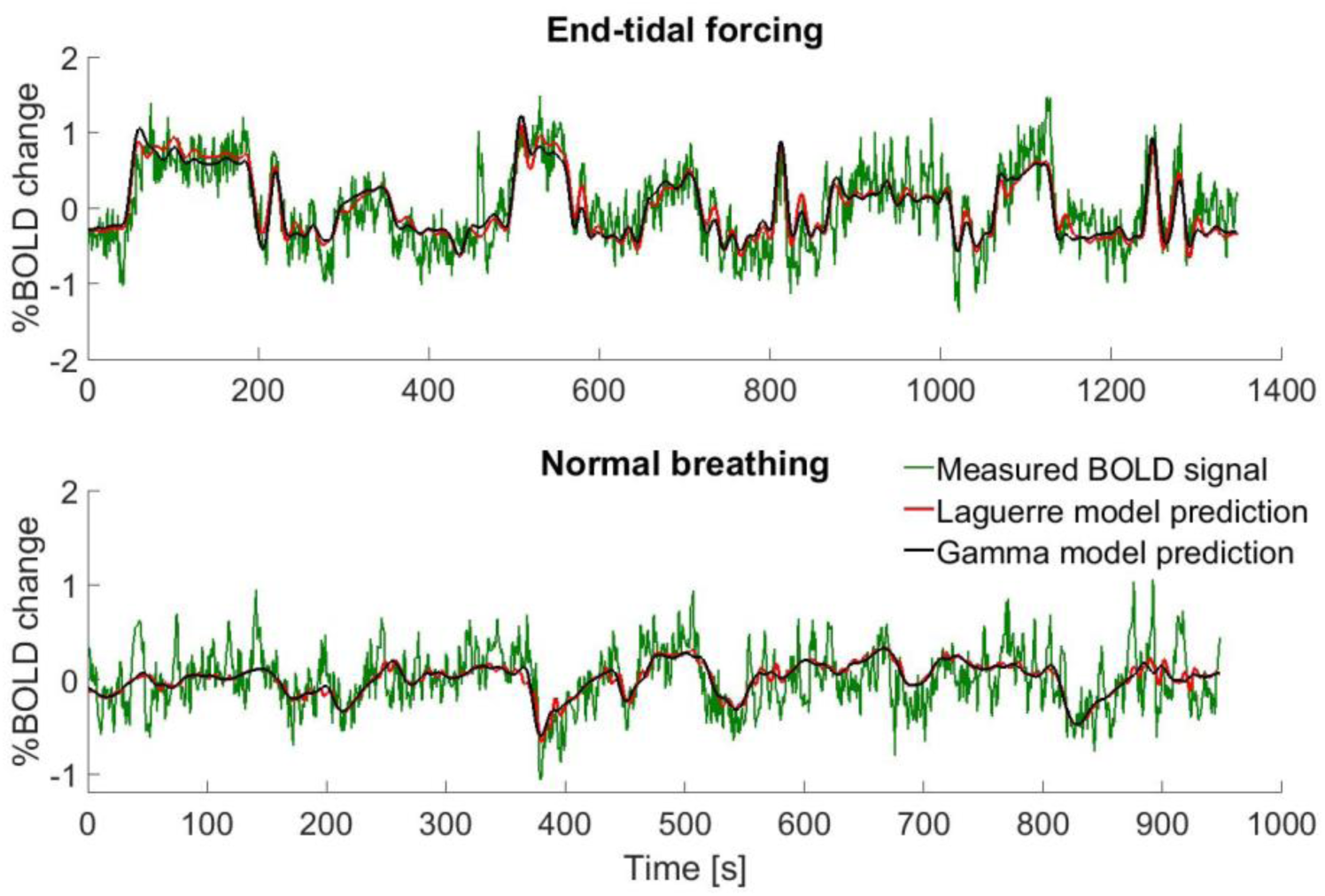
Representative gamma and Laguerre model output predictions for the left anteroventral nucleus of the thalamus functional ROI during end-tidal forcing (top panel) and resting breathing (bottom panel) conditions. The gamma and Laguerre models yielded similar model predictions, which explained a large fraction of the slow variations in the BOLD signal during end-tidal forcing and normal breathing (resting state conditions).

### 3.1.2. Voxel-wise analysis

Fig. 5 shows representative maps of voxel-specific features extracted from the corresponding dCVR curves, during externally induced hypercapnic CO2 challenges (end-tidal forcing) (left column) and normal breathing (right column). To increase the image contrast, each map was adjusted such that 1% of the lower and higher intensity values were saturated. The extracted features include area (steady-state CVR), time-to-peak (time of maximum instantaneous amplitude), absolute peak (maximum instantaneous amplitude), and power (sum-of-squares). The area maps (Fig. 5(a)) obtained under forcing conditions generally exhibit higher intensity values compared to the maps obtained under resting conditions (Fig. 5(b)). Subcortical structures such as the thalamus and the brainstem, as well as regions in the cerebral cortex show increased sensitivity to the CO2 challenges. In contrast, white matter shows lower sensitivity under both forcing and resting conditions. Under forcing conditions, periventricular white matter regions show exquisitely small steady-state CVR values. Similar patterns can be observed for the absolute peak (Fig. 5(e,f)) and power (Fig. 5(g,h)) maps. The time-to-peak maps (Fig. 5(c,d)), in which lower intensity values correspond to faster response time, show that for many regions in the brain the timing of the maximum instantaneous amplitude of dCVR is generally slower during forcing conditions.

**Fig. 5.**
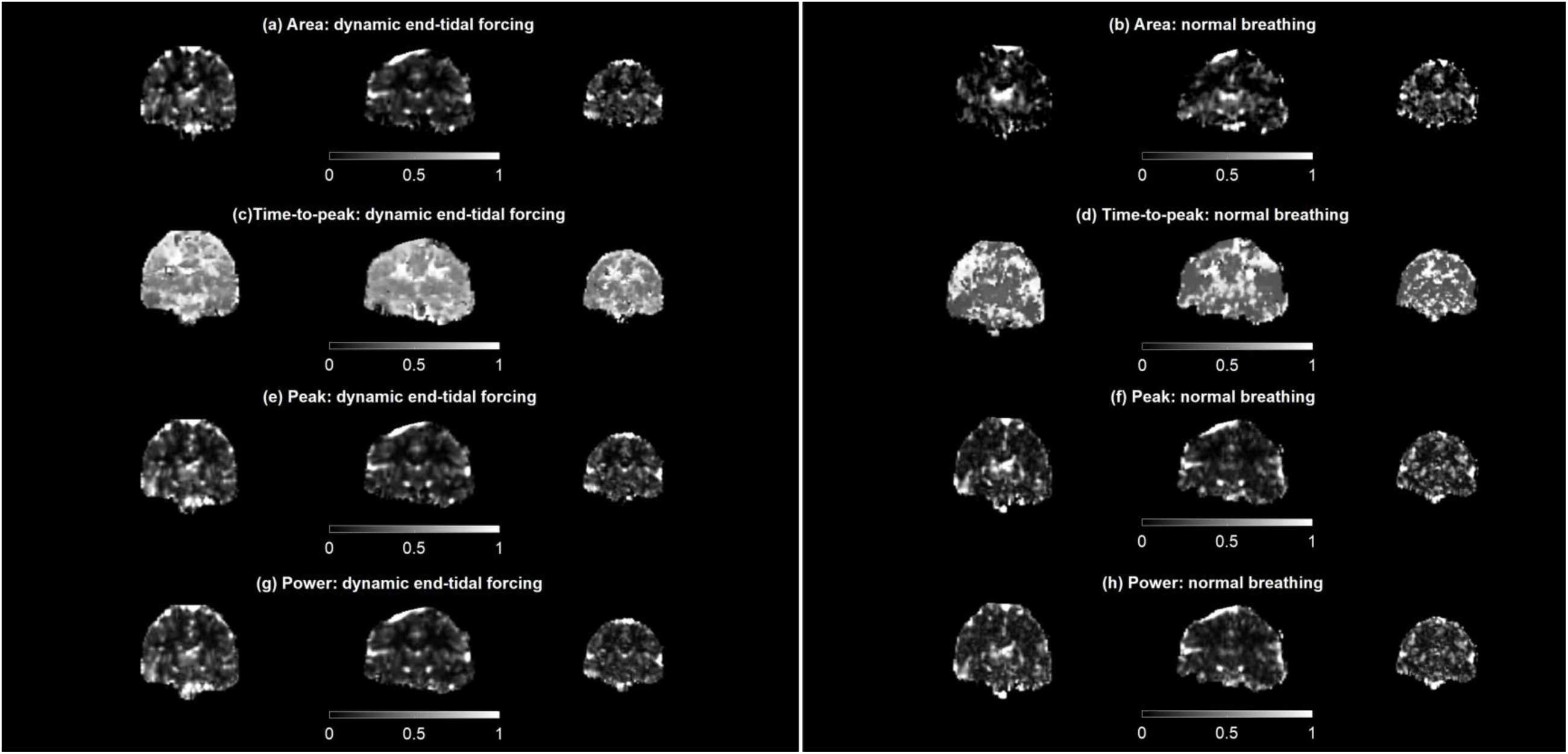
Regional maps of voxel-specific dCVR features from three representative subjects during end-tidal forcing (left column) and resting (right column) conditions. First row (a,b): dCVR area. This feature corresponds to the steady-state CVR value. Second row (c,d): dCVR time-to-peak. This feature corresponds to the time lag of the maximum instantaneous effect of CO2 on the BOLD signal. Third row (e,f): Peak dCVR value. This feature corresponds to the maximum instantaneous effect of CO2 on the BOLD signal. Fourth row (g,h): dCVR power values. This feature corresponds to the dCVR curve sum-of-squares. The area, peak and power maps exhibit similar patterns of feature variability across different brain regions, revealing increased sensitivity to CO2 challenges in areas such as the brainstem, thalamus and cerebral cortex. White matter is generally less sensitive to the CO2 challenges compared to gray matter, with periventricular white matter regions exhibiting the lowest sensitivity. The time-to-peak maps show that the timing of the maximum instantaneous peak value of dCVR is slower during forcing conditions, suggesting that CO2 reactivity to larger CO2 challenges is slower compared to spontaneous fluctuations.

Fig. 6 illustrates the results of the one-way, nonparametric statistical comparisons (permutation paired test (Winkler et al. 2014) between the dCVR feature maps obtained under end-tidal forcing conditions compared to the corresponding maps obtained under resting conditions. The comparisons were performed on FSL (Jenkinson et al. 2012), after registration of the individual feature maps to the MNI-standard space. *P*-values were corrected using the threshold-free cluster enhancement method (TFCE) (Smith and Nichols 2009).

**Fig. 6.**
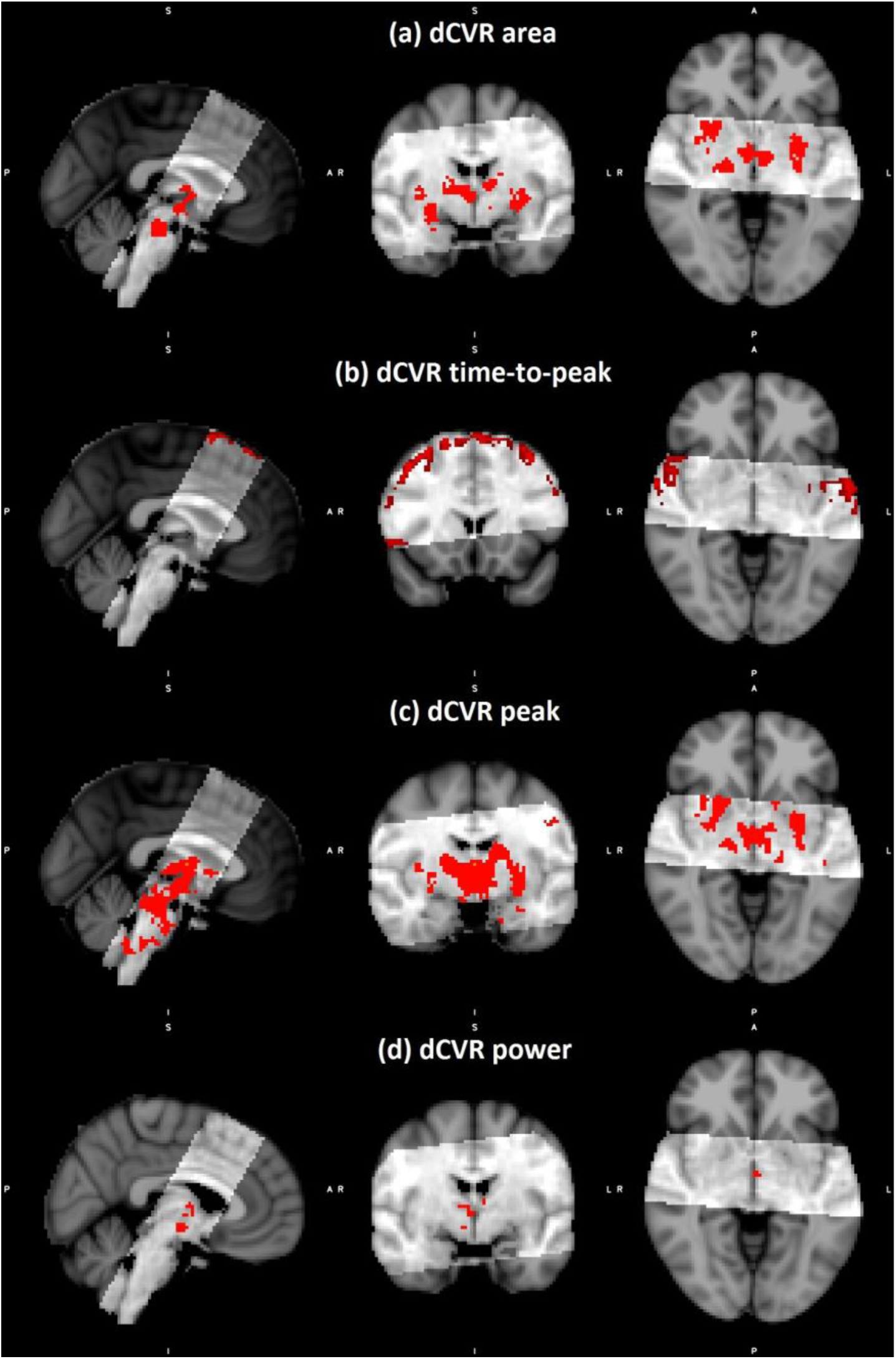
One-way comparisons of the dCVR feature values at each voxel between end-tidal forcing and resting conditions, after registration of the individual feature maps to the MNI standard space. In all cases the voxels corresponding to significantly different feature values (p<0.0005) are shown in red, whereby the p-values were corrected using the TFCE method (Smith and Nichols 2009). First row (a): dCVR area. Second row (b): dCVR time-to-peak. Third row (c): dCVR peak value. Fourth (bottom) row (d): dCVR power. The comparisons of the area, peak and time-to-peak maps show significant differences in the anterior nuclei, and the ventral posterior lateral nuclei of the thalamus. The comparisons of the area and peak maps revealed increased sensitivity in the pons and the putamen during forcing conditions. In contrast, the comparison of the time-to-peak maps revealed significant differences in cortical regions, including the insular, posterior frontal and premotor cortices.

The comparisons of the area, peak and time-to-peak maps obtained during CO2 challenges compared to resting fluctuations suggest significant differences in the anterior, and the ventral posterior lateral nuclei of the thalamus. The comparisons of the area and peak maps under the two conditions, respectively, also revealed significant differences in the pons, as well as in the putamen. In contrast, the comparison of the time-to-peak maps revealed significant differences in cortical regions, including the insular, posterior frontal and premotor cortices, suggesting that these areas reach their peak dCVR values. There were no areas yielding significantly larger values for all features (area, time-to-peak, peak, energy) during resting fluctuations compared to C02 challenges.

### 3.1.3. Clustering analysis

Table 2 illustrates the number of clusters that resulted from the classification analysis of the dCVR curve shapes using k-means clustering and the silhouette criterion for evaluating classification performance and selecting the number of clusters (Kaufman et al. 2005; Rousseeuw 1987). For all subjects except one, the optimal number of clusters varied between four and five. For subject 6, the optimal number of clusters during resting conditions was found to be six. Fig. 7 shows the mean dCVR curve of each cluster that resulted from the clustering analysis of voxel-specific dCVR curves obtained from a representative subject, for both experimental conditions. The cluster indices were selected so that mean dCVR curves that are overall more negative correspond to a smaller index values, whereas mean dCVR curves that are overall more positive correspond to greater index values.

**Table 2.** Total number of clusters of dCVR curve shapes for each subject. The clustering was performed using k-means clustering and the silhouette criterion for evaluating clustering performance and selecting the optimal number of clusters.

|  | Dynamic end-tidal forcing | Normal Breathing |
| --- | --- | --- |
| Subject 1 | 5 | 4 |
| Subject 2 | 4 | 4 |
| Subject 3 | 5 | 5 |
| Subject 4 | 5 | 5 |
| Subject 5 | 4 | 4 |
| Subject 6 | 5 | 5 |
| Subject 7 | 4 | 4 |
| Subject 8 | 4 | 4 |
| Subject 9 | 4 | 5 |
| Subject 10 | 5 | 6 |
| Subject 11 | 5 | 4 |
| Subject 12 | 5 | 4 |

**Fig. 7.**
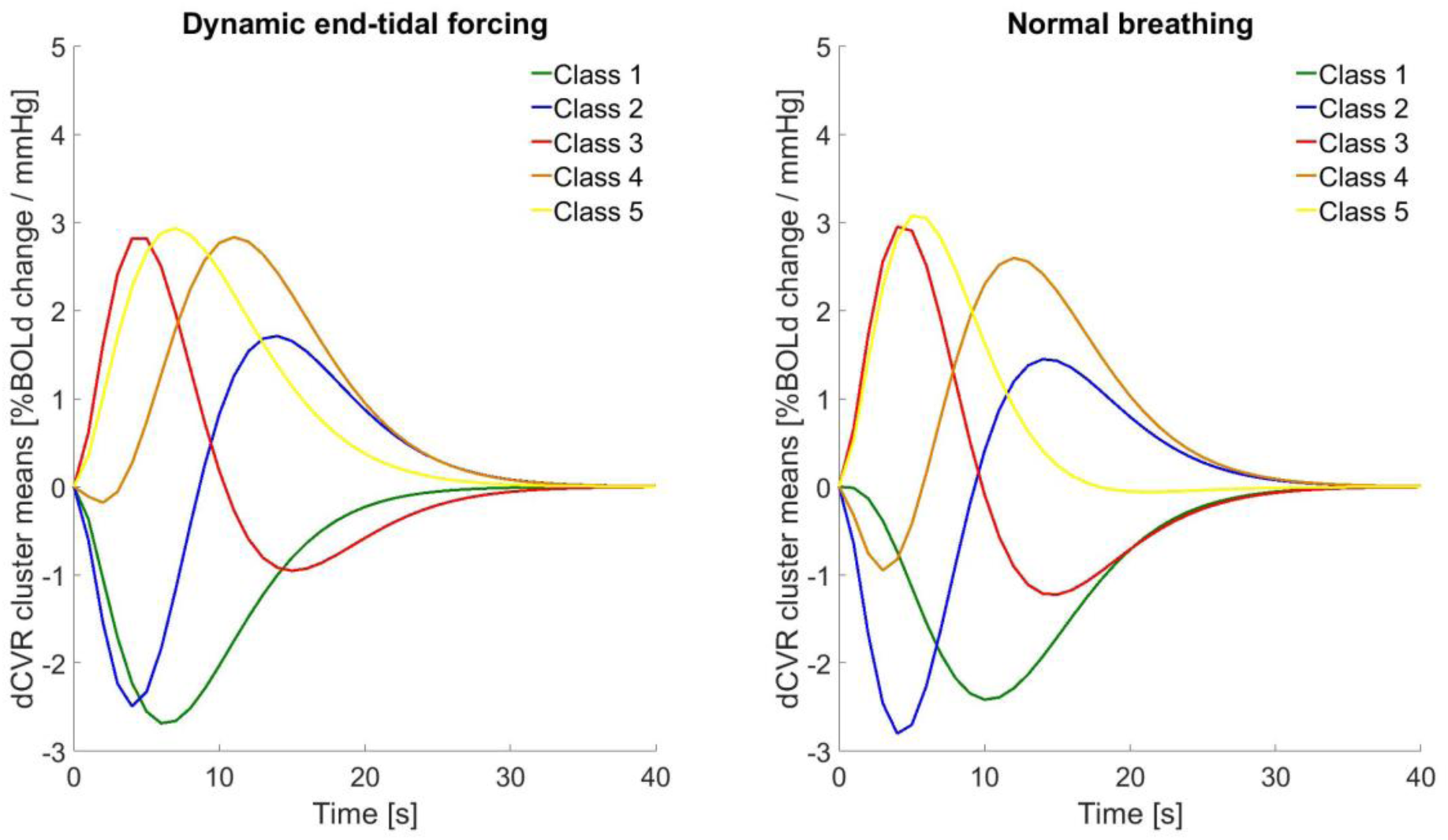
Mean dCVR curves of clusters obtained using k-means clustering and the silhouette criterion for evaluating classification performance and selecting the optimal number of classes of voxel-specific dCVR curves obtained from a representative subject. Left panel: End-tidal forcing. Right panel: normal breathing. The cluster indices were selected so that mean dCVR curves that are overall more negative correspond to a smaller index values, whereas mean dCVR curves that are overall more positive correspond to greater index values.

Figs. 8 illustrates the distribution of cluster indices within the GM, WM, CSF anatomical ROIs, within the brain volume of the same three representative subjects shown in Fig. 5, under externally induced hypercapnic CO2 challenges (end-tidal forcing). Smaller cluster index values represent more negative dCVR curve shapes, whereas higher index values represent more positive dCVR curve shapes. The histogram below each anatomical ROI map displays the distribution of the voxels inside that ROI into the clusters formed. Each histogram is normalized with respect to the total number of voxels within the ROI associated with it. In GM, the majority of voxel-specific dCVR responses to step CO2 challenges are classified into cluster 5. In WM, while most of the voxel-specific dCVR responses are classified into cluster 5, the number of voxels classified into cluster 2 is higher compared to GM. This suggests that WM has more voxels responding with an initial undershoot to step CO2 challenges compared to GM. This effect is more pronounced in CSF-rich regions, where, in comparison to GM, the number of voxel-specific dCVR curves classified into cluster 1 is lower and the number of voxel-specific dCVR curves classified into clusters 1-3 is higher.

**Fig. 8.**
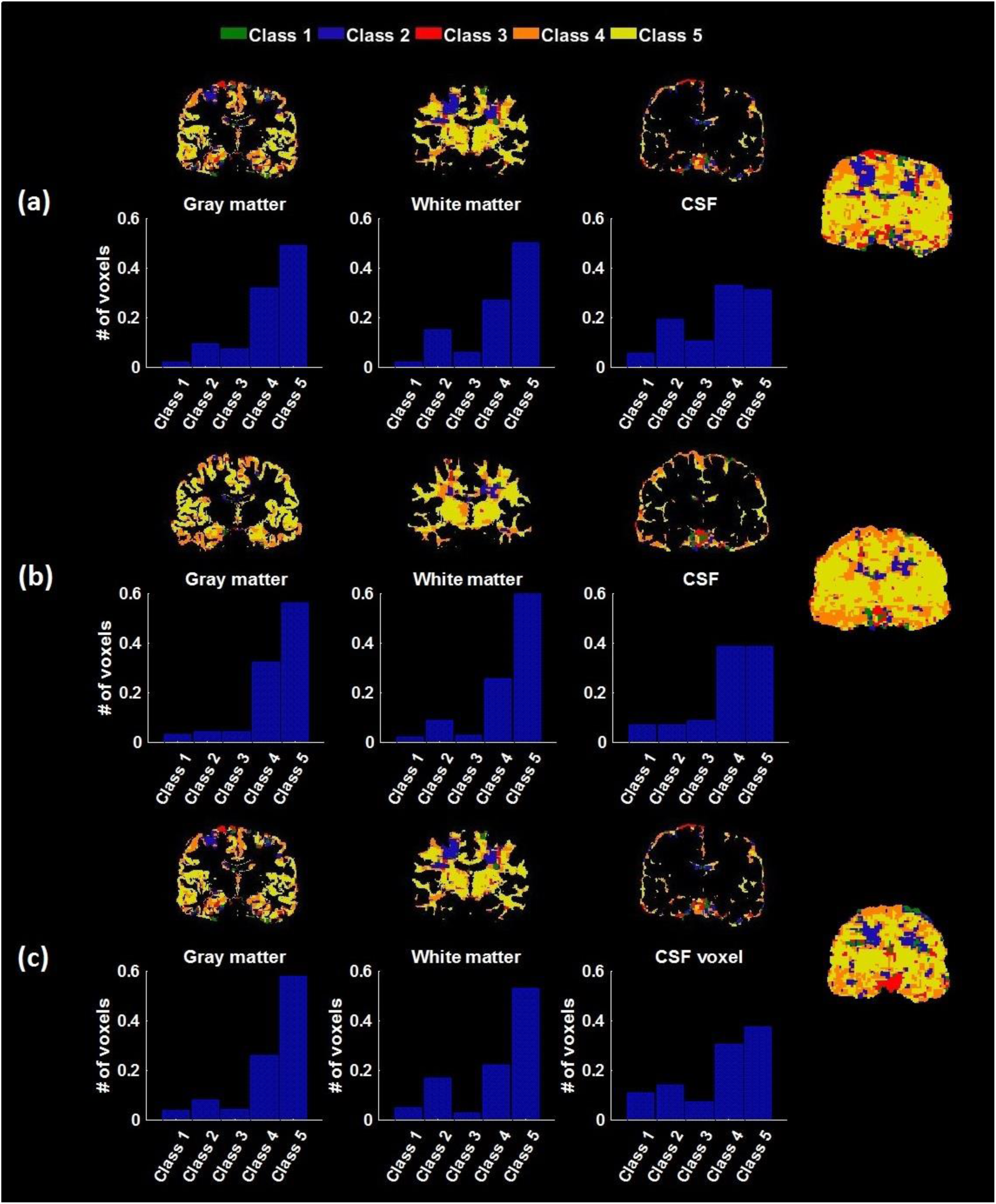
Representative maps of the cluster spatial distribution within the GM, WM, and CSF anatomical ROIs as well as the entire brain volume for 3 representative subjects during end-tidal forcing conditions. Smaller cluster index values correspond to more negative dCVR curve shapes, whereas higher index values correspond to more positive dCVR curve shapes. Representative dCVR cluster means are shown in Fig. 7(a). The histogram below each anatomical ROI map displays the distribution of ROI voxels into the clusters formed after application of the clustering analysis. The histograms were normalized with respect to the total number of voxels in each anatomical structure. The vast majority of voxels in GM are classified in cluster 5. Similarly, the majority of voxels in white matter are classified in cluster 5; however, the proportion of voxels classified into cluster 2 is increased compared to gray mater. In CSF regions, the proportion of voxels classified in cluster 5 is decreased compared to GM and WM, whereas the proportion of voxels classified in clusters 1-3 is increased.

Fig. 9 illustrates the distribution of cluster indices within the same anatomical ROIs and total brain volumes, as well as the histogram associated with each ROI, for the same subjects shown in Fig. 8, during normal breathing. In contrast to forcing conditions, where most of the GM voxel-specific dCVR responses are classified into cluster 5, the largest proportion of voxel-specific responses are classified into cluster 3, which corresponds to dCVR curve shapes characterized by an early overshoot followed by a late undershoot. This explains the different dCVR shapes for larger ROIs shown in Fig. 3.

**Fig. 9.**
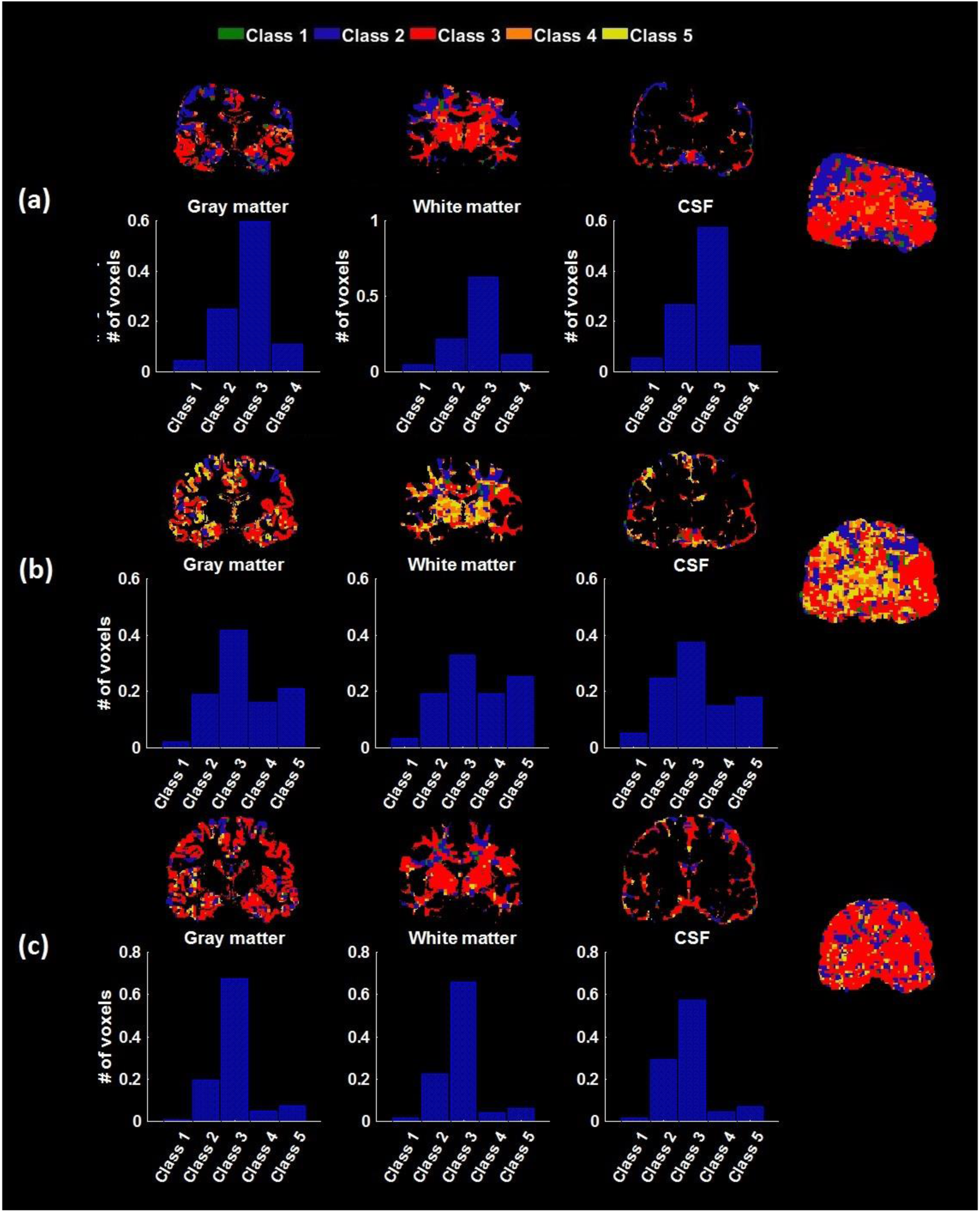
Representative maps of the cluster spatial distribution within the GM, WM, and CSF anatomical ROIs as well as the entire brain volume for 3 representative subjects during normal breathing (resting state) conditions. Smaller cluster index values correspond to more positive dCVR curve shapes, whereas higher index values correspond to more positive dCVR curve shapes. Representative dCVR cluster means are shown in Fig. 7(b). The histogram below each anatomical ROI map displays the distribution of ROI voxels into the clusters formed after application of the clustering analysis. The histograms were normalized with respect to the total number of voxels in each anatomical structure. The vast majority of voxels in all structures are classified into cluster 3, which corresponds to dCVR curve shapes characterized by an early overshoot followed by a late undershoot.

## 4. Discussion

We investigated the regional variability of dynamic cerebrovascular reactivity by modeling the dynamic interactions between CO2 and BOLD in healthy subjects during resting conditions and hypercapnic step changes induced by dynamic end-tidal forcing. To this end, we employed an efficient systems identification technique (functional expansions) to obtain estimates of dCVR curves within single voxels and larger ROIs, whereby we constructed a custom basis set by using the Laguerre and gamma basis sets (Section 2). Based on this, we demonstrated that dCVR exhibits significant regional variability that suggests the dynamic effect of CO2 on the BOLD signal strongly depends on brain region. Our results suggest that the proposed methodology is able to obtain robust dCVR estimates in single voxels even during resting conditions, despite the low SNR associated with the latter. This is supported by the similarity of the main dCVR features between resting and forcing conditions (Fig. 5), as well as by the similar cluster mean curve shapes, which resulted from the clustering analysis (Fig. 7).

This has important implications as it suggests that it is feasible to obtain reliable dCVR maps without the need of externally induced stimuli (end-tidal/prospective forcing, controlled breathing) which makes it easier to implement and applicable to a potentially wider class of patient populations. Our analysis revealed considerable variability of the dCVR estimates in both the large ROIs and individual voxels.

In the respiratory control centers of the brain (anteroventral thalamus, ventrolateral thalamus, ventral respiratory group, Kolliker-Fuse parabrachial group, ventral posterior lateral thalamus, retrotrapezoid nucleus – (Pattinson et al. 2009)), it was found that the estimated dCVR curves varied in their peak values and the timings of their peaks. Fig. 10 shows box plots of dCVR peak and time-to-peak values across all subjects obtained in the respiratory related centers of the brain, under forcing and resting conditions. The red vertical bars represent medians and the black circles mean values. The left and right anteroventral nuclei of the thalamus and the retrotrapezoid nucleus exhibit higher peak values compared to other ROIs, under both externally induced hypercapnic CO2 challenges (end-tidal forcing) and resting conditions. Under normal breathing conditions, the peak values of the dCVR estimates in the ROIs across all subjects are overall smaller compared with CO2 challenges. Moreover, under normal breathing conditions, the time-to-peak values of the dCVR estimates in the ROIs across all subjects are overall smaller compared to forcing conditions (*p* < 0.05; Kruskal-Wallis nonparametric one-way ANOVA test). In addition, the dCVR estimates during resting conditions exhibited a late undershoot that was absent during dynamic end-tidal forcing conditions (Fig. 3). These observations can be explained by the fact that under normal breathing conditions, the baseline level was maintained at 1 mmHg above the subjects’ natural P_ET_CO2, inducing hypercapnic vascular tension (Halani et al. 2015).

**Fig. 10.**
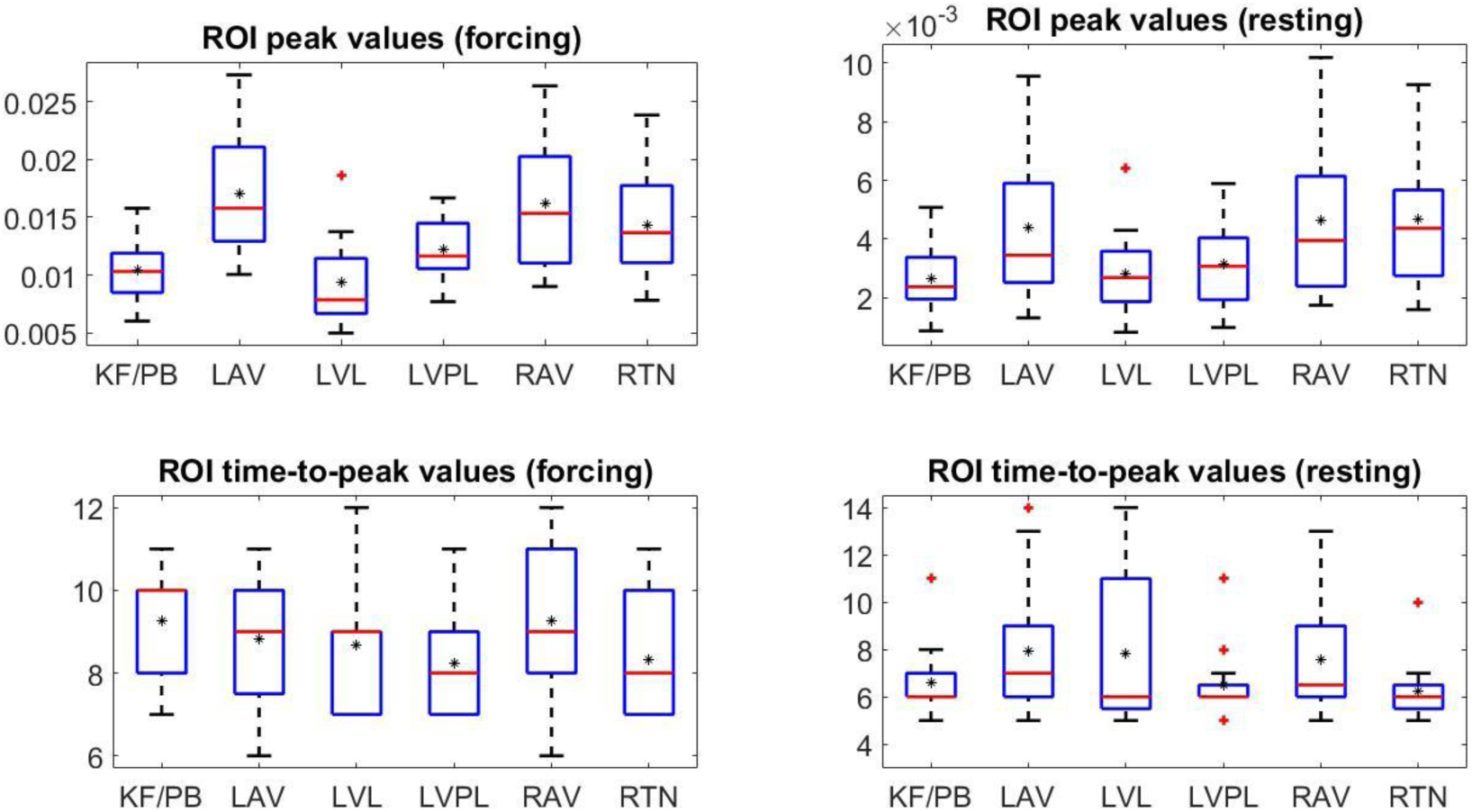
Box plots of dCVR peak and time-to-peak values across subjects obtained from the respiratory centers of the brain across all subjects (ROI analysis), under forcing (left column) and normal breathing (right column). The red vertical bars correspond to median values and the black circles correspond to mean values. The left and right anteroventral nuclei of the thalamus and the retrotrapezoid nucleus exhibit higher peak values compared to other ROIs, under both externally induced hypercapnic CO2 challenges and resting conditions. Under normal breathing conditions, the peak values of the dCVR estimates in the ROIs across all subjects are overall smaller compared to forcing conditions. Moreover, under normal breathing conditions, the time-to-peak values of the dCVR estimates in the ROIs across all subjects are overall smaller compared to forcing conditions (*p* < 0.05; Kruskal-Wallis nonparametric one-way ANOVA test). KF/PB: Kolliker-Fuse parabrachial group, LAV: left anteroventral thalamus, LVL: left ventrolateral thalamus, LVPL: left ventral posterior lateral thalamus, RAV: right anteroventral thalamus, RTN: Retrotrapezoid nucleus.

The individual maps of voxel-specific dCVR features, shown in Fig. 5, demonstrate the variability of the dCVR in a higher spatial resolution. The area, peak, and time to peak maps are similar to each other during both experimental conditions. The maps obtained under CO2 challenges have generally higher intensity values compared to normal breathing. This is more pronounced in the area maps due to the late undershoot in the dCVR curves obtained under normal breathing conditions, which is generally absent from the dCVR curves obtained under forcing conditions. In addition, the time-to-peak values during normal breathing conditions were found to be overall smaller compared to forcing conditions. This suggests that the response time of the vasculature to step CO2 challenges (forcing conditions) is generally larger (i.e. the peak instantaneous value is reached more slowly) than the response time to spontaneous CO2 fluctuations (normal breathing conditions).

The statistical comparisons of voxel-specific dCVR area, peak, and power values between resting and end-tidal forcing conditions reveal significant differences mostly in the anterior and ventral posterior lateral nuclei of the thalamus. Furthermore, voxel-specific dCVR area and peak values were found to be significantly different in the pons and the putamen. These findings are in general agreement with the voxel-wise results reported in (Pattinson et al. 2009), whereby it was found that the areas with the stronger response during external CO2 challenges were the bilateral anterior nuclei of the thalamus, the right posterior putamen, the left ventrolateral nuclei of the thalamus, and in midline in the dorsal rostral pons, the inferior ventral pons, and the dorsal and ventrolateral medulla.

The comparison of the time-to-peak values between the two experimental conditions reveals significant differences in cortical regions as opposed to subcortical regions. This suggests that cortical structures, consisting primarily of gray matter, respond faster to spontaneous fluctuations of P_ET_CO2 that occur during normal breathing compared to step-changes in P_ET_CO2 that occur under baseline hypercapnia. It is known that in the presence of a hypercapnic stimulus, there is a larger CBF increase in gray matter structures (especially cortical regions) compared to normocapnia (normal breathing), a fact shown both in humans and rodents (Ramsay et al. 1993; Sicard and Duong 2005). As a result, the vasculature in gray matter may require more time to adapt to the large changes in CBF that occur under forcing conditions in response to step changes in CO2 compared to smaller changes in CBF that occur in response to spontaneous fluctuations in P_ET_CO2, which explains the significant differences in the dCVR time-to-peak values observed in cortical regions of the brain.

It is worth noting that significant differences in BOLD occurring in response to externally induced CO2 step challenges compared to normal breathing that are observed in some brain regions, in addition to vascular reactivity, might be due to differences in neural activation. These neural activation - induced changes in BOLD seem to be (perhaps unsurprisingly) reflected on features depending on the entire shape of the dCVR curve, such as total area, rather than individual measures that only depend on instantaneous values of the dCVR curve, such as peak value. This is shown e.g. in Fig. 6(a) where the statistical comparisons of dCVR area values between resting and end-tidal forcing conditions reveal significant differences in the brainstem and the thalamus, including regions implicated in the generation and control of respiration rhythms (Pattinson et al. 2009). On the other hand, the corresponding dCVR peak value (Fig. 6(b)) and power (Fig. 6(d)) are more extensive compared to the voxel-wise maps presented in (Pattinson et al. 2009), as the area is the only measure that takes into account the sign of instantaneous CO2 effects. Finally, the dCVR time-to-peak maps (Fig. 6(c)) revealed significant differences in cortical regions, which were not identified in (Pattinson et al. 2009).

The clustering analysis of the voxel-wise dCVR estimates (Figs. 7-9) revealed that the dCVR shapes are distributed symmetrically across the brain, a result which was reproducible across most of the subjects. In the case of end-tidal forcing (Fig. 8), the largest part of the brain WM and GM was assigned to clusters which corresponded to voxel - specific dCVR curves with more positive curve shapes. On the other hand, in the case of normal breathing (Fig. 9) the number of dCVR curve shapes exhibiting a late undershoot in the brain WM and GM is overall larger compared to end-tidal forcing.

During hypercapnia, periventricular WM regions were found to exhibit negative steady-state CVR values (Fig. 5(a)). Similar results were also reported in (Mandell, Han, Poublanc, Crawley, Kassner, et al. 2008), where the authors attributed it to “vascular steal”. This can be seen more clearly from the clustering of the voxel-specific dCVR curves in the brain WM, shown in Fig. 11, where periventricular WM regions yielded dCVR curves that are classified into clusters characterized by prevalently negative dCVR curves (clusters 1 and 2) compared to the rest of the brain WM. Similarly, CSF-rich regions in the brain, such as the lateral ventricles, yielded a larger proportion of dCVR curves that were classified to clusters characterized by prevalently negative dCVR curves (clusters 1 and 2) compared to GM and WM, as it is shown by the histograms in Fig. 8. This agrees with the findings of (Thomas et al. 2013), where the ventricular BOLD signal was found to be anti-correlated with hypercapnic step changes in CO2, which was attributed to CSF movement due to the large blood volume increase that occurs in response to the large hypercapnic CO2 step changes.

**Fig. 11.**
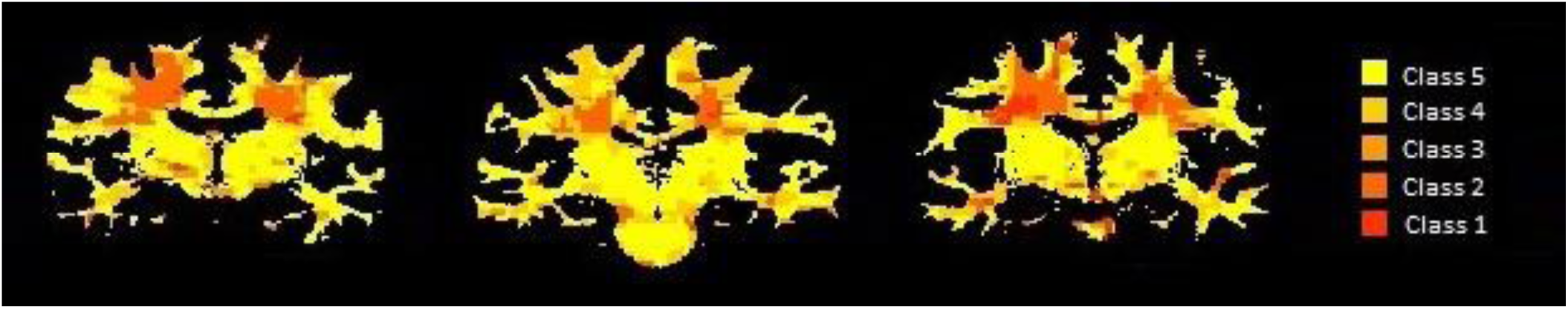
Representative results of the clustering analysis in white matter for three representative subjects (same subjects shown in Figs 8 and 9). While most of voxels are assigned to clusters 4 and 5 that are characterized by dCVR curves exhibiting a large overshoot, WM periventricular voxels are mainly classified in clusters with smaller indices, which are characterized by dCVR curves exhibiting a large initial undershoot.

During resting conditions, the clustering analysis revealed that voxel-specific dCVR curve shapes may be different compared to forcing conditions. In particular, Figs. 8 and 9 show that the histogram of the cluster indices in each anatomical structure changes between forcing and normal breathing conditions. This implies that the dCVR curve shape of a particular voxel might change in normocapnia with respect to hypercapnia, suggesting that the underlying mechanism of vasodilation might respond differently to CO2 fluctuations during normal breathing than to hypercapnic CO2 changes.

## 5. Conslusion

In this work, we used linear and non-linear models to investigative dynamic CO2 reactivity in the respiratory centers of the human brain during both resting breathing and hypercapnic externally induced step changes in CO2, using measurements from 12 healthy subjects. We concluded that in these regions dynamic CO2 reactivity is mainly linear, for both experimental conditions. Therefore, we rigorously investigated the regional variability of dynamic CO2 reactivity in individual voxels using linear models. In this context, we estimated voxel-specific dynamic CO2 reactivity curves, and we showed that the regional characteristics of these curves vary considerable across different brain regions, and that their shape might be different under the two experimental conditions. Finally, we performed clustering analysis on the shapes of the estimated curves, which resulted into clusters of similar curve shapes that were distributed symmetrically across the brain. Our results suggest that it is feasible to obtain reliable estimates of dynamic cerebrovascular reactivity curves from resting-state data, which could allow the design of safer and easier to implement clinical protocols for the assessment of dCVR, which do not require external stimuli (e.g. hypercapnia), in any patient population.

## Acknowledgements

KP is supported by the National Institute for Health Research Oxford Biomedical Research Centre based at Oxford University Hospitals NHS Trust and University of Oxford. KP has acted as a consultant for Nektar Therapeutics. The work for Nektar has no bearing on the contents of this manuscript.

